# Effects of Florivore Type and Plant Chemotype on Metabolic Responses in Flowers and Leaves of an Aromatic Plant Species

**DOI:** 10.64898/2026.11.13.766439

**Authors:** Selina Gaar, Lukas Brokate, Thomas Dussarrat, Caroline Müller

**Author notes:** **Correspondence:** Caroline Müller, Thomas Dussarrat.

## Abstract

Florivory is an important biotic challenge in plant-insect interactions, evoking induced responses in plant specialised metabolism, the extent of which depends on phenotypic plasticity. While responses to leaf herbivores are well studied, floral responses to florivores with different feeding modes, their systemic effects on leaves and the role of intraspecific chemodiversity remain poorly understood. We used clonal replicates of five *Tanacetum vulgare* (Asteraceae) chemotypes, which differ in their leaf and flower terpene profiles, to test how florivory by chewing beetles (*Olibrus* spp.) and phloem-feeding aphids (*Macrosiphoniella tanacetaria*) affects the metabolic responses of flower heads and systemically in leaves. Floral volatile organic compounds (VOCs) and stored terpenes were analysed via GC-MS and metabolic fingerprints via LC-MS. While overall metabolic profiles of flower heads and leaves remained largely stable following florivory, specific metabolites shifted in abundance locally and systemically. These changes primarily involved the shikimate and phenylpropanoid pathways and minor terpenes. The specific metabolic responses varied by florivore species and chemotype, a pattern observed across floral VOCs, as well as stored terpenes and untargeted metabolic fingerprints, differing also between organs. Overall, florivory triggers organ-specific chemical responses in reproductive and vegetative tissues, shaped by a complex interplay between plant chemotype and florivore type.

## 1. Introduction

Plants interact with a diverse community of insects that feed on leaves, roots, stems and/or flowers. Flowers, in particular, are often nutrient-rich resources for mutualists such as pollinators, making them also attractive targets for floral herbivores, known as florivores (McCall and Irwin, 2006; Sasidharan et al., 2023). As reproductive organs, flowers contribute strongly to fitness and according to the optimal defence hypothesis, plants should therefore allocate defences preferentially to them. Flowers are expected to be protected by high constitutive concentrations of specialised metabolites, whereas in leaves induced defences are often more pronounced (Brown et al., 2003; McCall and Irwin, 2006; Smallegange et al., 2007). Florivores differ in their feeding modes, which determine how and where they damage flowers and how plants respond. Biting-chewing florivores, such as beetles and caterpillars, consume floral tissue directly and often cause visible damage, whereas piercing-sucking florivores, such as aphids, feed on phloem sap, typically at the floral stalks (Rusman et al., 2019). Chewing mainly activates the jasmonate/ethylene pathway, while piercing-sucking typically activates the salicylic acid pathway, although crosstalk occurs (Erb et al., 2012; Koornneef and Pieterse, 2008; Schweiger et al., 2014). Plant responsiveness to feeding by insects with different feeding modes has mostly been studied in leaves (Mithöfer and Boland, 2012; Schuman and Baldwin, 2016), while knowledge on flower responses is comparatively scarce.

Signalling pathways shape downstream defence responses, including changes in specialised metabolite profiles (Walling, 2000). In flowers, florivory can alter profiles of both emitted volatiles and stored specialised metabolites (Lucas-Barbosa et al., 2016; Zangerl and Berenbaum, 2009). Altered emission of volatile organic compounds (VOCs) potentially acts as indirect defence by attracting natural enemies (Desurmont et al., 2015; Schiestl et al., 2014), whereas changes in the stored metabolite composition may contribute to direct defence against subsequent florivores (Boyer et al., 2016; Rusman et al., 2022). However, volatile and stored metabolites can also be attractive or stimulatory to plant antagonists (Gols, 2014; Schiestl, 2015). Flowers therefore face a trade-off: they must repel/deter florivores while remaining attractive to mutualists, which suggests that floral defences should be selective (Kessler and Halitschke, 2009; McCall and Irwin, 2006; Schiestl et al., 2014). Possible mechanisms include visitor-specific blends of attractive and repellent VOCs (Kessler et al., 2013; Raguso, 2016) or temporal regulation of VOC emission (Theis et al., 2007). Moreover, defence responses can also be systemic, as exemplified by the observed variation in stored and volatile floral metabolites under leaf herbivory (Bruinsma et al., 2014; Rusman et al., 2022; Schiestl et al., 2014). Because flowers and leaves are connected via the phloem, which determines how far systemic induction extends (Orians, 2005), signals from damaged flowers may also reach leaves through the vascular system. In accordance, florivory has been shown to alter leaf chemistry, such as glucosinolates in *Brassica nigra* (Brassicaceae) after combined florivore and pathogen attack (Chrétien et al., 2022) and anthocyanins in *Impatiens capensis* (Balsaminaceae) after artificial florivory (Soper Gorden and Adler, 2016). However, whether and how florivory affects leaf chemistry across plant species remains largely unknown.

Plant specialised metabolite profiles also differ constitutively among individuals of the same species in composition and concentration, a phenomenon referred to as intraspecific chemodiversity (Moore et al., 2014; Wetzel and Whitehead, 2020). In some species, this variation is structured into so-called chemotypes (Müller et al., 2026), i.e., groups of individuals that differ in the presence and/or quantity of their dominant metabolites, for example alkaloids or terpenoids in species of Asteraceae (Castells et al., 2014; Grof-Tisza et al., 2022). As a consequence, chemotypes also differ in their interactions with herbivores, affecting, for example, herbivore preference behaviour and performance (Jakobs and Müller, 2018; Kleine and Müller, 2011; Sasidharan et al., 2023; van Leur et al., 2008). Chemotypes may also differ in their inducibility, expressing phenotypic plasticity. For example, chemotype-specific differences in induced VOC emissions and stored metabolites were found in *Gossypium hirsutum* (Malvaceae) and *Tanacetum vulgare* (Asteraceae) after feeding by chewing and phloem-feeding herbivores (Clancy et al., 2020; Mamin et al., 2025; Newrzella et al., 2026). Despite the potential ecological relevance, the knowledge about induced chemotype-specific responses is still limited.

Among the Asteraceae, *T. vulgare* (common tansy) stands out for its pronounced intraspecific chemodiversity in leaf and floral terpenes across a broad geographical range (Clancy et al., 2016; Keskitalo et al., 2001; Wolf et al., 2011). Leaf terpene profiles allow individuals to be grouped into chemotypes, each defined by its dominant and minor co-occurring terpenes (Clancy et al., 2018; Kleine and Müller, 2011). Within its natural range, *T. vulgare* is colonised by a diverse herbivore community (Schmitz, 1998) and chemotype-specific plant responses have been demonstrated. For example, infestation by the specialised aphid *Macrosiphoniella tanacetaria*, which preferentially feeds on the stems below the inflorescence, alters the nutritional quality of the phloem sap, with effects depending on the colonised plant part and the plant’s chemotype (Jakobs et al., 2019). Infestation by this aphid species on leaves also enhanced shoot monoterpene emissions in a chemotype-specific manner (Newrzella et al., 2026). Flowers of *T. vulgare* are also visited by oligophagous *Olibrus* beetles (Coleoptera: Phalacridae), which feed mainly on pollen and other parts of flower heads (Eilers et al., 2021; Sasidharan et al., 2023). When feeding on different stems of the same plants under field conditions, *M. tanacetaria* and *Olibrus* spp. induced chemotype-specific metabolic responses in the flower heads: aphids mainly decreased and beetles mainly increased the levels of metabolic features, with little overlap in significant features across insect species and plant chemotypes (Gaar et al., 2026). Nevertheless, both insect species increased features related to the shikimate and phenylpropanoid pathways (Gaar et al., 2026). Yet, it remains unclear how chemotypes shape local and systemic plant responses to these distinct florivore types.

The aim of this study was to elucidate the effects of florivory by insects differing in feeding mode on the metabolic responses in both flower heads and leaves of different *T. vulgare* chemotypes. Plants of five chemotypes were cloned and clones were either infested with *M. tanacetaria* aphids or *Olibrus* spp. beetles or kept uninfested as controls, allowing us to test plant individual-specific responses. Metabolic responses were measured using a multi-platform approach, analysing floral VOCs as well as terpenes and metabolic fingerprints of flower heads and leaves to determine local as well as systemic responses. We expected plant responses to depend on the chemotype and florivore feeding mode, with only modest shifts in overall metabolic profiles restricted to individual metabolites in both flower heads and leaves. Specifically, we hypothesised florivory to cause an increase in individual floral VOCs, but only minor changes in stored flower and leaf terpenes. Moreover, we expected alterations across diverse other classes of specialised metabolites, especially those associated with the shikimate and phenylpropanoid pathways, with metabolites modulated by aphid infestation being predominantly decreased in abundance, whereas those affected by beetle feeding to be predominantly increased and more pronounced.

## 2 Materials and Methods

### 2.1 Plant and Insect Material

Seeds of different *T. vulgare* plants (considered as different maternal genotypes) were collected in 2019 in Bielefeld, Germany (for details see Ziaja and Müller, 2023). Plants were grown in a greenhouse and chemotypes assigned based on leaf monoterpene profiles. Five chemotypes were selected for further studies, based on their abundance and stability and to maximise chemical diversity. Two chemotypes were dominated by a single terpene accounting for >50% of the total terpene content, namely β-thujone (BThu) and artemisia ketone (Keto), while three chemotypes were characterised by two to three major terpenes contributing >50% in total, namely artemisyl acetate, artemisia ketone and artemisia alcohol (Aacet), α- and β-thujone (ABThu) and (*Z*)-myroxide, santolina triene and artemisyl acetate (Myrox). As *T. vulgare* is an outcrossing species, offspring from one maternal individual can belong to different chemotypes. To separate chemotype from maternal genotype effects, we chose plants from multiple maternal origins per chemotype. In total, the selected chemotypes originated from seeds of ten maternal genotypes, with overlapping genotypes across chemotypes where possible. From each plant individual, three clones were produced from rhizomes and cultivated in separate pots (0.5 L) in a 1:1 mixture of sterilised potting soil and river sand. Plants were grown in a climate chamber at 23 °C during the light and 18 °C during the dark period, with a relative humidity of 60–70%. The photoperiod was initially set to 12 h light/12 h dark for three weeks, increased to 14 h light/10 h dark for two weeks and then maintained at 16 h light/8 h dark for the rest of the experiment, intending to induce flowering. Nevertheless, plants originating from different individuals started flowering at different time points, but clones within plants were roughly synchronised (mean flowering-onset difference: 21.3 ± 25.4 days). The lights provided an average UV-A radiation of 0.524 W m⁻² and a photosynthetically active radiation of 180 μmol m⁻² s⁻¹. Plants were watered three times per week and positions rotated weekly. Once roots emerged through the pot bottom, plants were fertilised weekly with WUXAL Super 8-8-6 (Hauert MANNA Düngewerke GmbH, Germany) at a concentration of 2 mL L⁻¹ of water. *Neoseiulus cucumeris* were released preventively every eight weeks to control for potential thrips and *Steinernema feltiae* were applied every three months against fungus gnats.

A colony of *Macrosiphoniella tanacetaria* aphids was established in May 2024 from individuals collected in Bielefeld and maintained on *T. vulgare* plants grown from purchased seeds of a random mixture of chemotypes (as described in Gaar et al., 2026). Aphids were reared in tents (60 × 60 × 60 cm) in a climate chamber (16 h light/8 h dark; 22 °C/18 °C; 60-70% relative humidity). *Olibrus* spp. beetles (probably *Olibrus aeneus*) were collected in summer 2024 in Bielefeld and Salzkotten from flower heads of wild chamomile species (*Matricaria* spp., *Anthemis* spp., or *Tripleurospermum* spp., Asteraceae) and maintained in plastic boxes (22 × 13 × 12 cm) lined with paper tissue within the same climate chamber as the aphids (as described in Gaar et al., 2026). They were provided with flower heads of *T. vulgare* from a mixture of all five chemotypes used in this study (greenhouse-grown) as well as flower heads from wild *T. vulgare* plants.

### 2.2 Performance of Experiment and Sampling

At peak flowering, each of the three clones was either infested with *M. tanacetaria*, with *Olibrus* spp. or kept uninfested as control (Figure 1; Table S1). To maintain insects on flower heads, polyethylenetetraphtalate sauce cups (135 mL, Pro-Pac Ostendorf Plastic, Germany) with removable lids and mesh bags were used. A 6 cm diameter hole was cut into the bottom of each cup and a mesh bag with its lower end cut open was glued to the cup outside. The cup was slipped over a flower head bouquet and fixed in position to a stick. Depending on the bouquet size, 5-15 flower heads were enclosed, two apterous adult aphids per flower head or one adult beetle per three flower heads were placed into the cup to standardise florivory pressure and the bag was closed around the stem with a drawstring ribbon. The infestation levels reflect natural florivore densities observed on *T. vulgare*.

**FIGURE 1.**
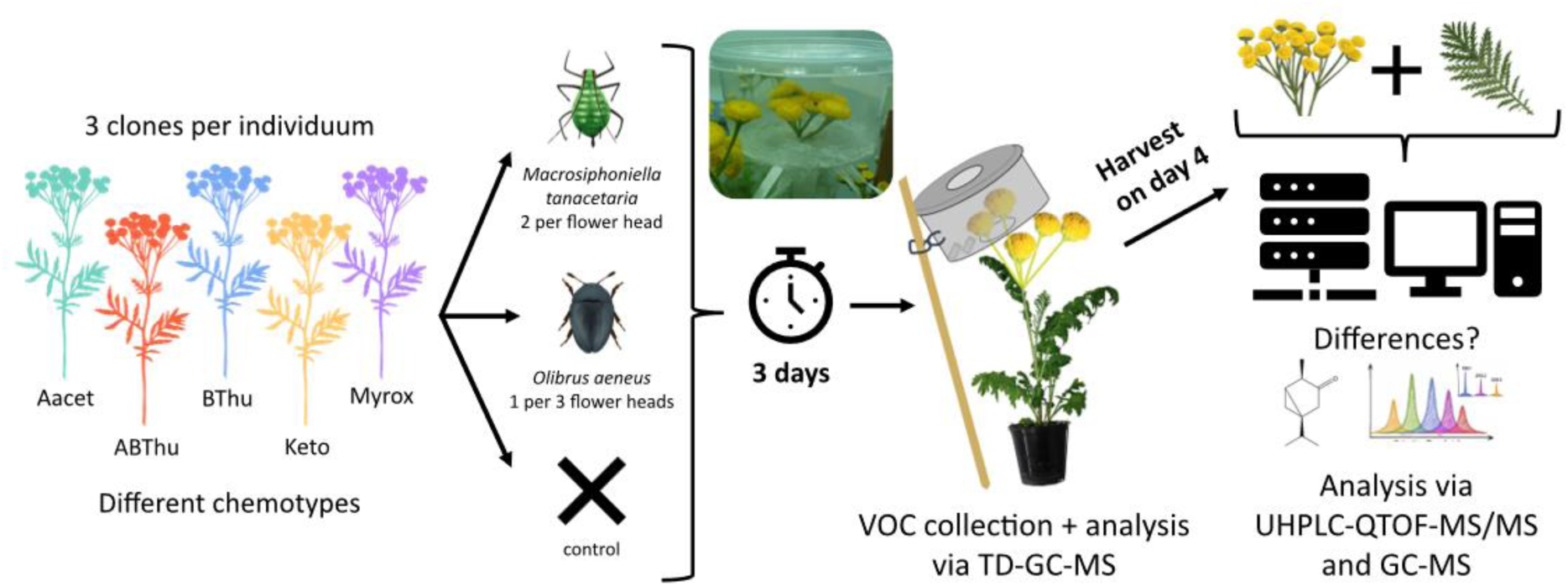
Experimental setup for testing chemotype-specific flower and leaf responses of *Tanacetum vulgare* to infestation by florivorous insects. Plants of five chemotypes [Aacet, artemisyl acetate/artemisia ketone/artemisia alcohol mix chemotype: *n* = 6; ABThu, α-/β-thujone mix chemotype: *n* = 7; BThu, β-thujone chemotype: *n* = 9; Keto, artemisia ketone chemotype: *n* = 4; Myrox, (*Z*)-myroxide/santolina-triene/artemisyl acetate mix chemotype: *n* = 4] were each cloned three times and each clone received one of three treatments (aphids, beetles, control). Depending on the number of flower heads per bouquet, 5-15 flower heads were enclosed in cups and either two aphids (*Macrosiphoniella tanacetaria*) added per flower head, one beetle (*Olibrus* spp.) per three flower heads or no insects (control). After three days, volatile organic compounds were collected on PDMS tubes (analysis via TD-GC-MS). On day four, flower heads and the second youngest unfolded leaf were harvested and analysed for terpenes (GC–MS) and metabolic fingerprints (UHPLC–QTOF–MS/MS).

Clones of the control treatment received empty cups in the same manner.

After three days of treatment, emitted volatiles were passively collected from late morning to afternoon, using polydimethylsiloxane (PDMS) tubes as sorbent material, modified after Eckert et al., (2023). Two PDMS tubes (Carl Roth, Karlsruhe, Germany) with an inner diameter of 1.0 mm, an outer diameter of 1.8 mm and a length of 0.5 cm were placed into the cups, avoiding any contact with the flower heads. To sample background volatiles of the climate chamber, one additional cup (hereafter cup blank) with two PDMS tubes per sampling event was fixed on a stick at similar height as the flower heads and positioned in a pot filled with the same substrate. After 6 h collection, the PDMS tubes were removed, placed in glass vials (1.5 mL), sealed with polytetrafluoroethylene tape and stored at −70 °C until chemical analysis. These VOC profiles may also contain VOCs released from the insects, which nevertheless contribute to the overall bouquet that a pollinator or other insect would encounter.

After four days of infestation, the insects were removed. Flower heads and the second youngest unfolded leaf were harvested, immediately flash-frozen in liquid nitrogen and stored at −70°C for later analysis of terpenes and metabolic fingerprints. Infestation and harvest were always conducted between 10 am and 12 pm. Since flowering started asynchronously among plants, 24 sampling events were conducted over a period of eight months. As some plants ultimately did not flower, the final number of successfully sampled triples was for Aacet: *n* = 6, ABThu: *n* = 7, BThu: *n* = 9, Keto: *n* = 4, Myrox: *n* = 4. In addition, for three Myrox individuals, only one of the three clones flowered, two of these clones treated with beetles, the other with aphids. For one additional Myrox individual, aphid- and beetle-infested clones were accidentally swapped during leaf sampling, so only the control leaf was retained for further processing. Prior to extraction, frozen flower heads and leaves were freeze-dried for 48 h and ground to a fine powder. The workflow is depicted in Figure 1.

### 2.3 Analysis of Emitted Volatiles and Stored Terpenes

Volatiles collected on PDMS tubes were analysed using a thermal desorption-gas chromatography-mass spectrometry (TD-GC-MS) system (TD-30 Series, GC 2010plus – MS QP2020, Shimadzu, Japan) operated in electron impact ionisation mode at 70 eV (similar as in Eckert et al., 2023; Ziaja and Müller, 2025). Thermal desorption of the tubes was carried out at 250 °C for 8 min with a flow rate of 60 mL min⁻¹. The cryogenic trap was held at −20 °C during desorption and subsequently heated to 280 °C for 3 min for analyte release. Injection into the GC was performed at 280 °C in splitless mode, with a helium column flow of 1.54 mL min⁻¹. Compounds were separated on a semi-polar VF-5 MS capillary column (30 m × 0.25 mm × 0.25 µm, 10 m guard column; Agilent Technologies, CA, USA). The oven temperature was initially set to 50 °C and held for 5 min after injection, then increased to 250 °C at a rate of 10 °C min⁻¹, followed by a ramp of 30 °C min⁻¹ to 280 °C, with a final hold for 2 min. The MS was operated with an ion source temperature of 230 °C, an interface temperature of 280 °C and a detector voltage of 1.5 kV. Mass spectra were acquired in quadrupole MS mode using the following scan parameters: from 8.2 to 19 min, a scan speed of 2000 and an *m*/*z* range of 30–400; from 19 to 29 min, a scan speed of 625 and an *m*/*z* range of 30–500. To calculate retention indices (RI) according to van den Dool and Kratz (1963), an alkane standard mixture (C7–C40; Sigma-Aldrich) was analysed by desorbing 1 µL from a PDMS tube using the same GC method. To prevent potential carry-over, an idle run was performed after the alkane standard and after every twelfth sample.

Stored terpenes were analysed following Eilers et al. (2021) with minor modifications. For each sample, 10 ± 2 mg were extracted in 1 mL *n*-heptane (HPLC grade, Roth, Germany) containing 10 ng µL⁻¹ 1-bromodecane (GC-MS grade, Sigma Aldrich, Germany) as internal standard, homogenised (FastPrep-24^TM^ 5G, MP Biomedicals, Germany) for 30 s at 9 m s⁻¹, sonicated in an ultrasonic bath for 5 min at room temperature and centrifuged for 10 min at 20 °C. The supernatants were collected for GC–MS analysis and blanks containing only the internal standard prepared in the same way (injected every 20 plant samples). Samples were analysed using the same GC–MS system as for the VOC measurements, but with liquid injection at 230 °C in split mode (1:10), a final hold time at the maximum temperature of the oven of 3 min and an MS interface temperature of 250 °C. An alkane standard mixture was analysed once a day, using the same GC–MS method to calculate RIs following van den Dool and Kratz (1963). Samples were analysed in two batches: the first after approximately three months of sampling events and the second after all plants had been sampled.

Chromatograms of VOCs and stored terpenes were processed using LabSolutions GC-MS Postrun Analysis v4.45 (Shimadzu). Compounds were identified by comparing their RI and mass spectra with those reported in Adams (2007), as well as with entries from the mass spectral libraries of NIST (2014) and FFNSC 3 (Mondello, 2015). A compound was considered identified when its mass spectrum matched a reference spectrum and its RI deviated by no more than 1.5% from the reported value (Adams, 2007), as previously described in Gaar et al. (2026). Remaining features were annotated as “unknown monoterpene or sesquiterpene”, based on characteristic masses. Compounds were quantified based on peak areas from the total ion chromatogram. For VOCs, the peak area of the cup blank collected on the corresponding sampling event was subtracted from the sample peak area and the values were normalised to the number of flower heads per sample. In approximately 80% of the VOC samples, acetophenone was overloaded and therefore excluded from further analyses. For stored terpenes, peak areas were normalised to the internal standard, mean peak areas of blanks were subtracted from each sample and values were subsequently normalised to the sample dry mass. Compounds detected in fewer than three samples were removed from the final table (Tables S2, S3, S4, S5).

### 2.4 Analysis of Metabolic Fingerprints

Metabolic fingerprinting was performed following Dussarrat et al. (2023) and Schweiger et al. (2021), with minor adaptations. For each sample, 10 ± 2 mg was extracted in 500 µL of 90% methanol (v/v) with hydrocortisone (10 mg L⁻¹; Sigma-Aldrich) as internal standard, vortexed, sonicated for 20 min in an ice-cooled ultrasonic bath and centrifuged for 10 min at 6 °C. The supernatants were filtered (0.2 µm, Phenomenex, Torrance, CA, USA) before LC–MS analysis. Extraction blanks containing the internal standard were prepared in the same manner (injected every 20 samples) and pooled quality control samples (QC) were prepared from aliquots of all extracts (injected every 15 samples).

Liquid chromatography-mass spectrometry (LC–MS) analyses were performed using a Dionex UltiMate 3000 UHPLC (Thermo Fisher Scientific, CA, USA) coupled to a compact quadrupole time-of-flight (QTOF)-MS/MS (Bruker Daltonics, Germany), equipped with a Kinetex XB-C18 column (150×2.1 mm, 1.7 µm, with guard column; Phenomenex), following Schweiger et al. (2021) with minor modifications. Chromatographic separation was carried out at 45 °C with a flow rate of 0.5 mL min⁻¹, using Millipore water with 0.1% formic acid as eluent A and acetonitrile with 0.1% formic acid as eluent B. The gradient began at 2% B, increasing to 30% over 20 min and then to 75% over 9 min, followed by column cleaning and re-equilibration. Mass spectrometric data were acquired in negative electrospray ionisation mode over an *m*/*z* range of 50–1300 at an acquisition rate of 5 Hz. All other instrument parameters followed Schweiger et al. (2021). MS/MS were acquired using AutoMS/MS mode. The isolation widths and collision energies were increased in line with the rising *m*/*z* of the precursor ions. Nitrogen served as the collision gas. Before each sample run, a sodium formate calibration solution was introduced to recalibrate the *m*/*z* axis.

Feature detection, alignment and grouping were carried out with the T-ReX 3D algorithm in MetaboScape (v. 2021b, Bruker Daltonics). Pre-processing of raw LC-MS data were performed as follows: intensity threshold of 1000 counts, minimum peak length of 11 spectra and EIC correlation threshold of 0.8. Features were retained only if eluted between 0.3 and 29 min and present in at least three samples. Recursive peak picking was applied with a minimum peak length of 6 spectra. Feature intensities were first normalised to the internal standard (hydrocortisone, [M+HCOOH−H]⁻). The coefficient of variation (CV) of each feature was then calculated across the QC samples and only stable features with a CV below 30% were retained. The remaining intensities were normalised to sample dry weight and the mean normalised blank intensities were subtracted from each corresponding feature, with negative values set to zero. The resulting datasets are provided in Tables S6 and S7.

### 2.5 Statistical Data Analysis

All data analyses and visualisations were performed in R v4.6.1 (R Core Team, 2026). Data were processed and plotted using the *tidyverse* collection of packages (Wickham et al., 2019), including *dplyr*, *tidyr*, *stringr*, *tibble*, *readr*, *purrr* and *ggplot2*. Heatmaps were created with *pheatmap* (Kolde, 2025), UpSet plots with *ComplexUpset* (Krassowski et al., 2022) and Venn diagrams with *VennDiagram* (Chen and Boutros, 2011).

To explore the overall variation in floral VOC profiles and to assess how plants of different leaf chemotypes cluster based on those, a principal component analysis (PCA) was performed on the normalised VOC data [relative values, cube root transformation and Pareto scaling performed in MetaboAnalyst v6.0 (Pang et al., 2024)]. To visualise how floral VOCs differ between treatments within each chemotype, a heatmap of the mean normalised abundance of each VOC was generated for every chemotype × treatment combination. To identify individual VOCs that responded significantly to florivory, moderated t-tests and fold changes (FC) were calculated for treatments *versus* controls, both across all samples and within each chemotype. Moderated t-tests were performed on the normalised data using the *limma* package (Ritchie et al., 2015), with plant identity included as a blocking factor to account for the paired design with plant clones. Moderated t-tests borrow information across all volatiles to stabilise variance estimates, which yields more reliable *P*-values when sample sizes are small, which was the case for the Keto and Myrox chemotypes. Only VOCs present in at least three samples were tested. Fold changes (FCs) were calculated from absolute abundances as the ratio of the insect treatment mean to the control treatment mean. *P*-values from the moderated t-tests were adjusted for multiple testing using the false discovery rate (FDR) method (Benjamini and Hochberg, 1995). As no VOC remained significant after FDR correction, VOCs were considered significantly increased or decreased based on unadjusted *p*-values (*p* < 0.05) combined with an FC > 2 or < 0.5. These VOCs were visualised in a heatmap based on normalised data. To compare responses across groups (chemotype × treatment combination), the overlap of significant VOCs, irrespective of the direction of change, was visualised in an UpSet plot.

The total stored terpene content (summed peak areas of all terpenes, normalised to internal standard and sample weight) was compared between flower heads and leaves within untreated control plants using a two-sided paired Wilcoxon signed-rank test. To explore variation in stored terpene profiles while accounting for the paired design with plant clones, a multilevel PCA was performed on the normalised data (relative values, cube root transformation and Pareto scaling performed in MetaboAnalyst) using the package *mixOmics* (Rohart et al., 2017). Multilevel PCA centres each terpene within plant clones, so that the remaining variation reflects clone differences. To visualise how the stored terpenes in flower heads and leaves differ among the treatment groups within each chemotype, a heatmap was generated per organ, showing the mean normalised abundance of each terpene. To identify insect species- and chemotype-specific modulations, moderated t-tests and FCs were calculated using the same approach as for the VOC data, separately for flower heads and leaves. Significant terpenes were visualised in a heatmap per organ and the overlap across groups (treatment × organ × chemotype combination) was visualised in an UpSet plot.

Variation in metabolic fingerprints was assessed by multilevel PCA and, separately for flower heads and leaves, by multilevel Partial Least Squares Discriminant Analysis (PLS-DA) on normalised LC-MS data using *mixOmics* (median normalisation, cube root transformation and Pareto scaling performed in MetaboAnalyst). To illustrate how metabolic profiles differ between flower heads and leaves across chemotypes, a heatmap was created for each organ, showing the mean normalised abundance of each feature per chemotype. Moderated t-tests and FCs were calculated as described for the VOCs. After FDR correction, only one of approximately 1400 features remained significant.

Chemical features were considered significantly altered if unadjusted *p*-values < 0.05 combined with an FC > 2 or < 0.5. To compare responses across groups, overlaps of significant features, irrespective of the direction of change, were visualised in UpSet plots per organ (chemotype × treatment) and across organs and treatments (chemotypes pooled). To compare the direction of responses, the percentage of increased features (the remainder, up to 100%, being decreased features) was shown in a bar plot for each treatment × organ × chemotype combination. To test whether the number of significant features differed among chemotypes, organs and insect species, counts of significant features per treatment × organ × chemotype combination (*n* = 20: 5 chemotypes × 2 organs × 2 insect species) were analysed. As the counts were overdispersed (dispersion of a Poisson model = 4.2), a negative binomial generalised linear model (GLM) was fitted using the MASS package (Venables and Ripley, 2002). The model included chemotype, insect species and organ as main effects and the interactions chemotype × insect and insect × organ. The proportion of increased *versus* decreased features (number of increased features out of all significant features) was analysed with a binomial GLM using the same predictors (dispersion = 1.1). Effects were tested with Type II likelihood-ratio tests (*car* package; Fox and Weisberg, 2019). When effects were significant, post hoc pairwise comparisons were performed on estimated marginal means with Tukey adjustment (package *emmeans*; Lenth et al., 2026). Chemical class annotation of significant features was performed using SIRIUS v6.5.4 (Dührkop et al., 2019). Full parameter settings are provided in Table S8. The metabolite structure and chemical class predictions were performed using CSI:FingerID (Hoffmann et al., 2022) and CANOPUS (Dührkop et al., 2021). For each feature, pathway, superclass and class, annotations in the Natural Products Classifier (NPC) ontology were retained when the confidence score was ≥ 0.8.

Significant features were assigned identification-confidence levels according to the Metabolomics Standards Initiative (MSI) guidelines (Schymanski et al., 2014), with MSI 2 obtained if cos score ≥ 0.7. For each pathway, the number of significantly increased and decreased features was counted across four comparison groups (aphid/beetle × flower heads/leaves) and visualised in a bar plot, regardless of the chemotype. Pie charts were plotted to show the distribution of significant features across chemical pathways for each chemotype.

To compare florivore-induced responses across the three platforms (floral VOCs, stored terpenes and metabolic fingerprints), bar plots were generated for each dataset showing the number of significant features that were specific to one chemotype or shared by two or more chemotypes, irrespective of insect species. For the same datasets, Venn diagrams (package *VennDiagram*; Chen and Boutros, 2011) were plotted to depict the overlap of significant features between aphid and beetle treatments, irrespective of chemotype. For stored terpenes and metabolic fingerprints, additional Venn diagrams showing the overlap between flower heads and leaves, irrespective of chemotype and insect species, were generated.

## 3. Results

### 3.1 Effects of Chemotype and Florivory on Floral VOC Emissions

In total, 136 putatively annotated floral VOCs were detected, which grouped according to chemotype in a standard PCA (Figure S1). The thujone-related chemotypes (ABThu, BThu) clustered distinctly from the Aacet, Keto and Myrox chemotypes, which share artemisia alcohol in their profiles. Aacet and Keto additionally share artemisia ketone, while Aacet and Myrox share artemisyl acetate. The overall VOC profiles did not separate by florivory treatment. However, the heatmap showed visible differences in the abundance of several VOCs between treatments within each chemotype (Figure S2). Analyses of all samples combined (i.e. considering all chemotypes) and chemotype-specific analyses identified a total of 30 VOCs that were significantly increased or decreased by florivory (*p* < 0.05, FC > 2 or < 0.5; Figure 2A; for detailed results of all moderated t-tests and FCs see Table S9). Responding volatiles were mostly specific to both the chemotype and the florivore species and only a few were shared among chemotypes or between the two insect species (Figure 2A, B). Aphid and beetle infestation led to a similar number of responsive VOCs overall, but this number varied among chemotypes depending on the insect species. Towards aphids, ABThu showed the strongest response (8 VOCs), whereas Myrox and BThu showed the weakest (1 VOC each). Towards beetles, Aacet showed the strongest response (5 VOCs), followed by Myrox and ABThu (4 each), whereas Keto and BThu had only one significant VOC change each.

**FIGURE 2.**
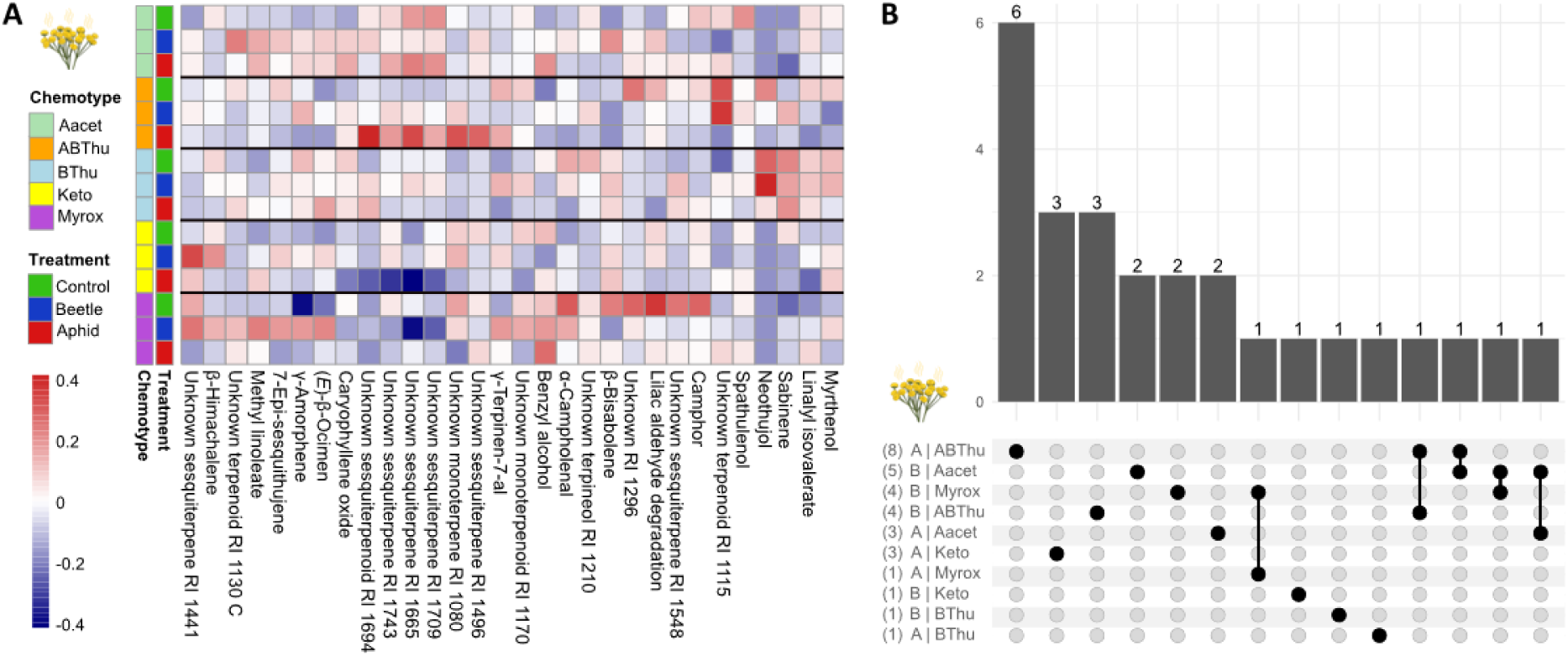
Changes of floral volatile organic compounds (VOCs) of *Tanacetum vulgare* of different chemotypes in response to florivory. Chemotypes are abbreviated as follows: Aacet, artemisyl acetate/artemisia ketone/artemisia alcohol mix chemotype; ABThu, α-/β-thujone mix chemotype; BThu, β-thujone chemotype; Keto, artemisia ketone chemotype; and Myrox, (*Z*)-myroxide/santolina-triene/artemisyl acetate mix chemotype. (A) Heatmap of the group mean values of normalised abundances of floral VOCs that differed significantly between treatments in moderated t-tests and fold change analyses (aphid or beetle versus control, *p* < 0.05; fold change < 0.5 for decreases or > 2 for increases), analysed across all samples and chemotype-specifically. Rows correspond to treatment groups within each chemotype, columns correspond to the significantly responding VOCs, clustered based on Pearson correlation and colours of different intensities represent normalised abundances. (B) Upset plot showing the number of significantly responding floral VOCs identified by moderated t-tests and fold changes, conducted for each chemotype individually. Numbers in parentheses on the left indicate the total number of significant VOCs within each group, while bars on the top represent intersections between groups, illustrating shared and unique volatile responses to insect infestation. In total, 136 floral VOCs were detected across all samples.

### 3.2 Effects of Chemotype and Florivory on Stored Terpenes in Flower Heads and Leaves

A total of 159 stored terpenes were detected in the flower heads and 149 in the leaves. All leaf terpenes were also present in the flower heads. The total stored terpene content was significantly higher in flower heads (median 427.6 a.u.) than in leaves (median 288.4 a.u.; *p* = 0.0013). Multilevel PCA revealed only a slight separation of flower head versus leaf profiles along PC1 (Figure 3A). Within organs, the stored terpenes showed treatment-related variations in their abundance within each chemotype (Figure S3). Analyses of all samples (i.e. considering all chemotypes) and chemotype-specific analyses identified a total of 20 stored terpenes that were significantly increased or decreased by florivory in flower heads and 12 in leaves (Figure 3B, C; for detailed results of all moderated t-tests and FCs see Table S10). In 8 of the 20 chemotype × insect × organ combinations, no terpene significantly responded to florivory. While mostly minor terpenes responded to florivory, the dominant terpene artemisia ketone decreased significantly in the flower heads of the Keto chemotype following aphid infestation. By contrast, in the ABThu chemotype, where artemisia ketone is not prominent, its abundance in flower heads increased significantly following both aphid and beetle infestation. Responding terpenes were largely specific to the organ, the chemotype and florivore species and only a few were significantly changed in both organs, multiple chemotypes, or by both insect species (Figure 3B-D). In flower heads, both insects affected a similar number of terpenes. In leaves, however, beetles led to a higher number of responding terpenes than aphids. The number of responsive terpenes also varied among chemotypes when organ and insect species were not considered, with the strongest response in Aacet (11), followed by ABThu (6), Myrox (4) and Keto (3) and the weakest in BThu (2).

**FIGURE 3.**
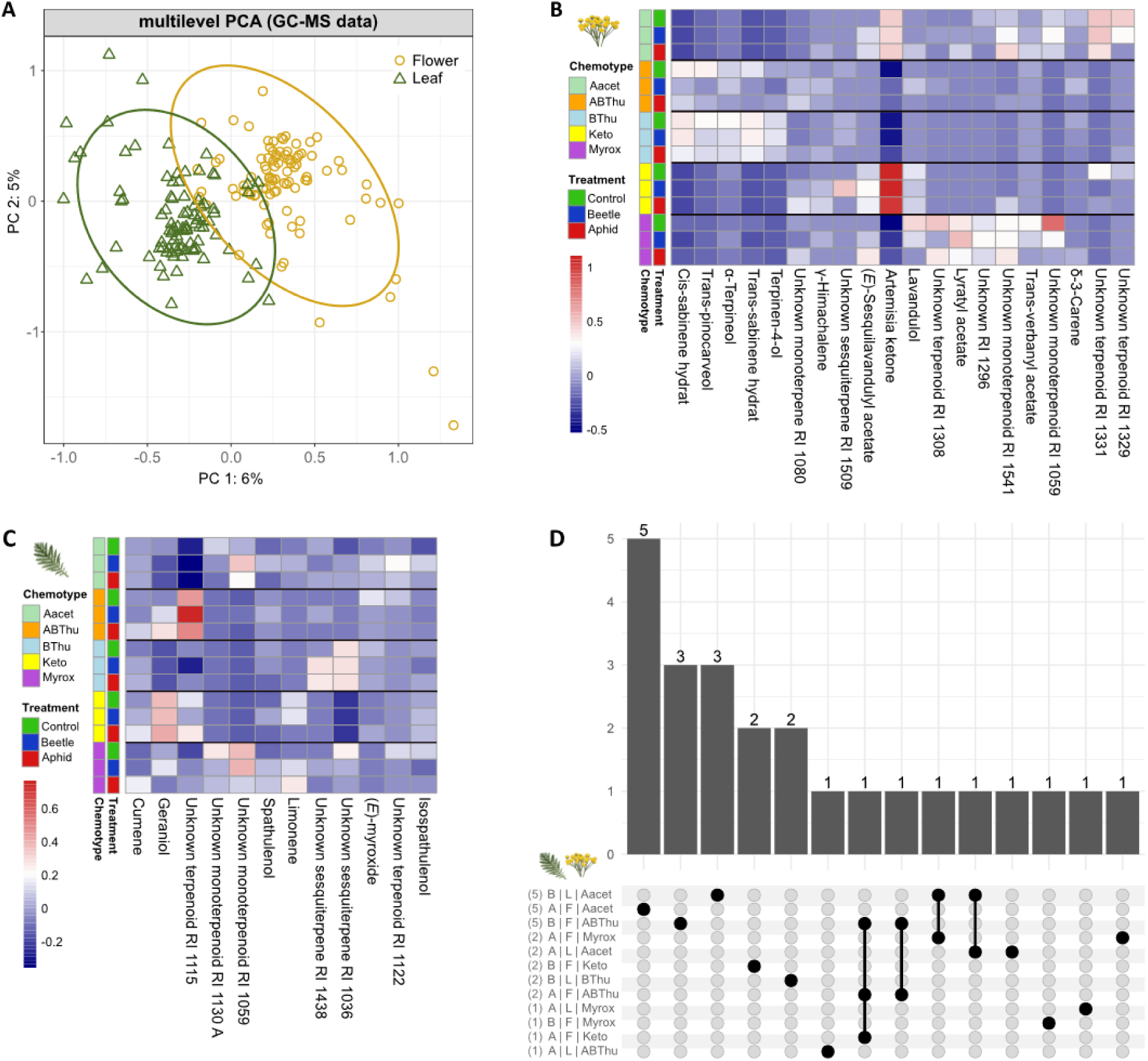
Stored terpene profiles of *Tanacetum vulgare* flower heads and leaves in response to florivory. (A) Multilevel PCA score plot showing the separation of samples between the two organs in a paired dataset comprising plant clones. Analysis was performed on normalised GC-MS data (159 terpenes). Ellipses indicate the 95 % confidence intervals for each group. (B, C) Heatmaps of the group mean values of normalised abundances of terpenes that differed significantly between treatments in moderated t-tests and fold change analyses (aphid or beetle *versus* control, *p* < 0.05; fold change < 0.5 for decreases or > 2 for increases) in (B) flower heads and (C) leaves. Analyses were performed using the pooled data set and chemotype-specifically. Rows correspond to treatment groups within each chemotype, columns correspond to the terpenes, clustered based on Pearson correlation and colours of different intensities represent normalised abundances. (D) Upset plot showing the numbers of terpenes significantly responding to florivory, identified by moderated t-tests and fold-change analyses for each organ-chemotype combination, aphid or beetle *versus* control. Numbers in parentheses on the left indicate the total number of significantly responding terpenes within each group, while bars on the top represent intersections between groups, illustrating shared and unique terpene responses to insect infestation. Only groups with at least one significantly responding terpene are shown; groups without significant responses were omitted from the plot. Abbreviations used in (B)-(D): Aacet, artemisyl acetate/artemisia ketone/artemisia alcohol mix chemotype; ABThu, α-/β-thujone mix chemotype; BThu, β-thujone chemotype; Keto, artemisia ketone chemotype; and Myrox, (*Z*)-myroxide/santolina-triene/artemisyl acetate mix chemotype; A, aphid; B, beetle; F, flower head; L, leaf.

### 3.3 Effects of Chemotype and Florivory on Metabolic Fingerprints in Flower Heads and Leaves

A total of 7163 features were detected in the flower heads and 7134 in the leaves, of which 7122 were shared. However, the abundance of many features differed substantially between the two organs (Figure S4). Several features differed also in abundance between chemotypes. Overall, multilevel PCA showed a clear separation between the metabolic profile of flower heads and leaves along PC1, which explained 55% of the variance (Figure 4A). PC2 explained only 4% of the variance and reflected a separation between individual plants, with all clones of a plant always clustering together. This separation was not related to the chemotype, treatment or plant age and its biological basis remains unclear. However, the supervised multilevel PLS-DA, performed separately for flower heads and leaves, revealed some separation of metabolic profiles among florivory treatments. In flower heads, the profiles of control samples were separated from those of insect-treated samples, whereas beetle- and aphid-treated samples slightly overlapped (Figure S5A). The control group was classified best by the first component alone, whereas the second component was needed to separate aphid- and beetle-treated samples. With two components, class error rates ranged from 0.31 to 0.39, indicating detectable but not strong treatment effects. In contrast to flower heads, the control group was not clearly separated in the multilevel PLS-DA for the leaves and the second component mainly improved the classification of the beetle-treated samples (Figure S5B). With two components, class error rates ranged from 0.41 to 0.46, indicating only weak separation.

**FIGURE 4.**
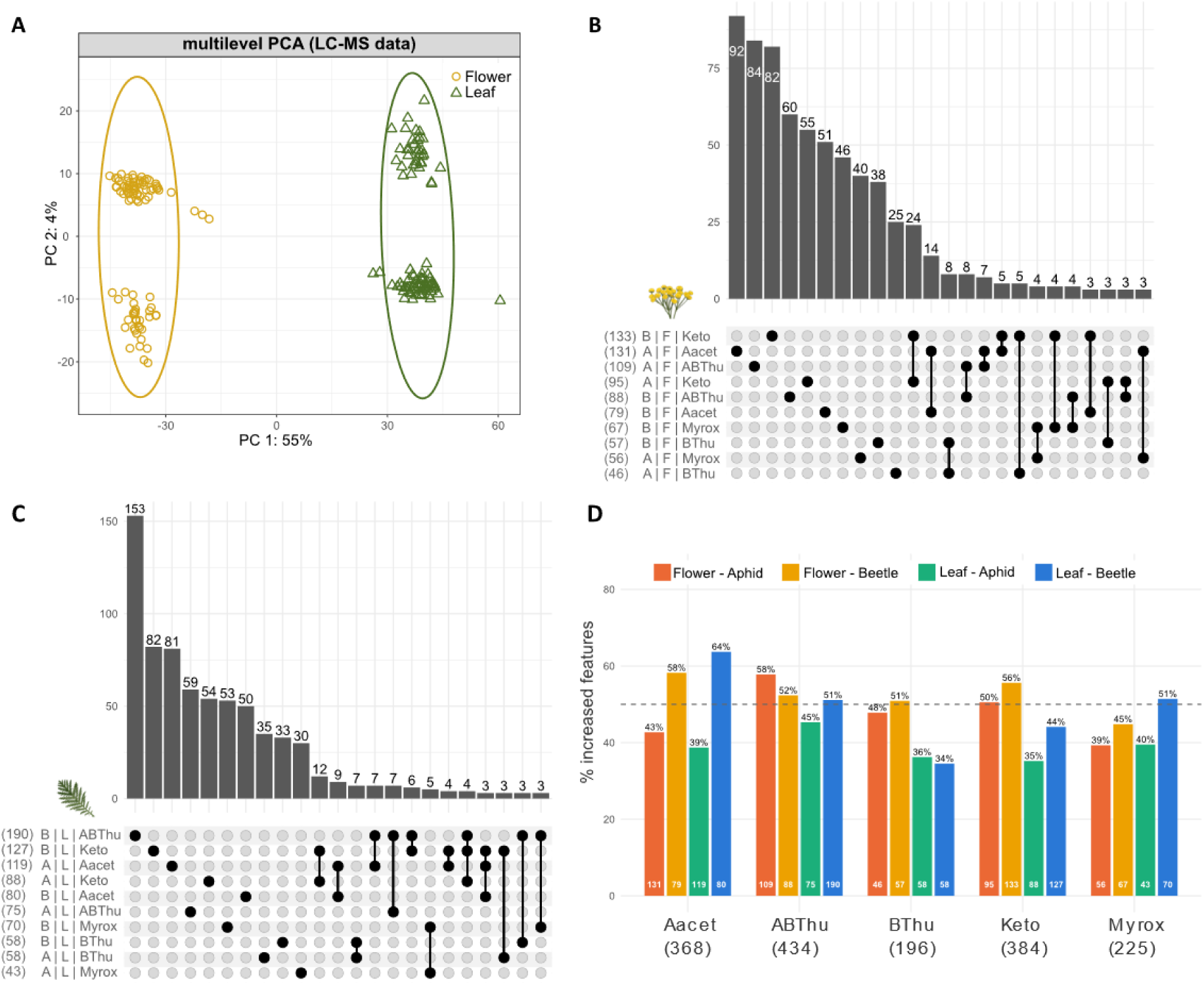
Metabolic fingerprints of *Tanacetum vulgare* flower heads and leaves in response to florivory. (A) Multilevel PCA score plot showing the separation of samples between the two organs in a paired dataset comprising plant clones. Analysis was based on normalised data of LC-MS metabolic fingerprints (7176 features). Ellipses indicate the 95 % confidence intervals for each group. Significant florivory-responsive features in (B-D) were identified by moderated t-tests and fold-change analyses conducted separately for each chemotype (aphid or beetle *versus* control, *p* < 0.05 and fold change < 0.5 for decreases or > 2 for increases). (B, C) Upset plots showing the numbers of features significantly responding to florivory in (B) flowers or (C) leaves. Numbers in parentheses on the left indicate the total number of significant features within each group, whereas top bars show intersections between groups, illustrating shared and unique metabolic responses to insect infestation. Only intersections of ≥ 3 features are shown. (D) Bar plot showing the percentage of significantly responding features with increased abundance relative to the control following florivory across chemotypes. Percentages are based on all significant features per chemotype, organ and treatment (total numbers are shown in white within the bars); the remaining percentage represents significantly decreased features. Numbers in parentheses below each chemotype indicate unique features per chemotype. The dashed horizontal line indicates 50 %, with values above or below indicating a predominance of increased or decreased features, respectively. Aacet, artemisyl acetate/artemisia ketone/artemisia alcohol mix chemotype; ABThu, α-/β-thujone mix chemotype; BThu, β-thujone chemotype; Keto, artemisia ketone chemotype; and Myrox, (*Z*)-myroxide/santolina-triene/artemisyl acetate mix chemotype; A, aphid; B, beetle; F, flower head; L, leaf.

Considering all chemotypes, 93 features were significantly increased or decreased by florivory across flower heads and leaves. There was no overlap between the features being changed in the two organs and only minimal overlap between features affected by the two insect species (*p* < 0.05, FC > 2 or < 0.5; Figure S6). In contrast, analyses performed separately for each chemotype identified a total of 1355 significantly modulated features, illustrating the need to consider intraspecific chemodiversity (for detailed results of all moderated t-tests and FCs see Table S11). Of these, 710 were significantly increased or decreased in flower heads and 756 in leaves, with 111 features being shared. Responding features were largely specific not only to the organ and chemotype, but also to the florivore species. Only a few significant features overlapped between insect species and chemotypes in both flower heads and leaves (Figure 4B, C).

Across all chemotype × insect × organ combinations, the number of significant features and the proportion of increased *versus* decreased features varied considerably (Figure 4D). The number of responding features varied significantly among chemotypes depending on the insect species (Table 1A). Under aphid feeding, Aacet showed more than twice as many significantly modified features (estimated mean: 125; Table S12) than BThu (52; *p* < 0.001) and Myrox (49; *p* < 0.001; Table S13), with ABThu (92) and Keto (91) having intermediate responses (significantly higher than in BThu and Myrox, all *p* < 0.01). In contrast, under beetle feeding, ABThu (137) and Keto (130) responded most, with both having more features significantly modified than Aacet (79; *p* < 0.001 and *p* < 0.01, respectively), BThu (57) and Myrox (68; all *p* < 0.001), the latter three not differing in responsiveness from each other (Tables S12, S13). Also, organ-specific effects depended on the insect species (Table 1). Beetles modulated a significantly higher number of features in leaves (98) than in flower heads (81; *p* < 0.05), whereas aphids elicited similar responses in both organs (82 in flower heads and 72 in leaves; *p* > 0.05; Table S14). However, the higher number of responding features in leaves under beetle feeding appeared to be restricted to ABThu (Figure 4D). The organ effect in beetles should therefore be interpreted with caution, as it may be driven by this chemotype. The proportion of increased *versus* decreased features depended on the interaction between chemotype and insect species and differed between organs (Table 1B). Only in the Aacet chemotype, beetle feeding caused mostly increased features (0.61), whereas aphid feeding was mainly associated with decreased features (0.59; odds ratio = 0.44, *p* < 0.001; Table S15). Across chemotypes and florivore type, increased and decreased features were balanced in flower heads (0.50 increased), whereas leaves showed a higher proportion of decreased features (0.56; Table S16), especially under aphid feeding (Figure 4D). Overall, more chemotype × insect × organ combinations showed mainly decreased than increased features (Figure 4D), although this trend was not tested statistically.

**TABLE 1:** Results of likelihood ratio tests (LR χ²) from the negative binomial generalised linear model for the number of responding features (A) and from the binomial generalised linear model for the proportion of increased features (number of increased out of all significant features, B), as affected by chemotype, insect species, organ and interactions.

| **Factor** | **LR χ²** | **df** | ***p*-value** |
|---|---|---|---|
| **A: Number of responding features** |  |  |  |
| Chemotype | 92.989 | 4 | **< 0.001** |
| Insect | 5.004 | 1 | **0.025** |
| Organ | 0.305 | 1 | 0.581 |
| Chemotype × Insect | 26.333 | 4 | **< 0.001** |
| Organ × Insect | 5.882 | 1 | **0.015** |
| **B: Proportion of increased features** |  |  |  |
| Chemotype | 7.494 | 4 | 0.112 |
| Insect | 10.636 | 1 | **0.001** |
| Organ | 6.639 | 1 | **0.010** |
| Chemotype × Insect | 9.981 | 4 | **0.041** |
| Organ × Insect | 1.211 | 1 | 0.271 |

LR χ² = likelihood ratio chi-square statistic; df = degrees of freedom; Significant effects (*p* < 0.05) are shown in bold

Of the 1399 features that responded significantly to florivory (1355 from the analyses performed separately for each chemotype and 44 additional features from the chemotype-independent analyses; Table S17, Figure 5), 15 could be annotated at the compound level (MSI 2), of which most belonged to the shikimate and phenylpropanoid pathways, including caffeic acid derivatives, flavonoids and other phenolic compounds, as well as an amino acid and a lipid (Table S17). The remaining features were classified at the level of biosynthetic pathway, superclass or class (Table S17). Pathway classification revealed that the shikimate and phenylpropanoid pathways responded most strongly to florivory (Figure 5A). In these pathways, aphid feeding affected more features in flower heads than in leaves, whereas beetle feeding showed the opposite pattern. Increased and decreased features were balanced in all four groups. Amino acids and peptides responded with similar numbers of features across insects and organs, but differed in direction: flower heads showed mainly increased features in response to both insect species, whereas leaves showed mainly decreased features. All other responding pathway classes (alkaloids, carbohydrates, fatty acids, polyketides and terpenoids) were mostly comparable between insects and organs in both the number of responding features and the proportion of increased and decreased features. Despite modest differences between chemotypes, florivory-responsive features showed broadly similar proportional distributions across chemical pathways when organ and insect species were not considered (Figure 5B).

**FIGURE 5.**
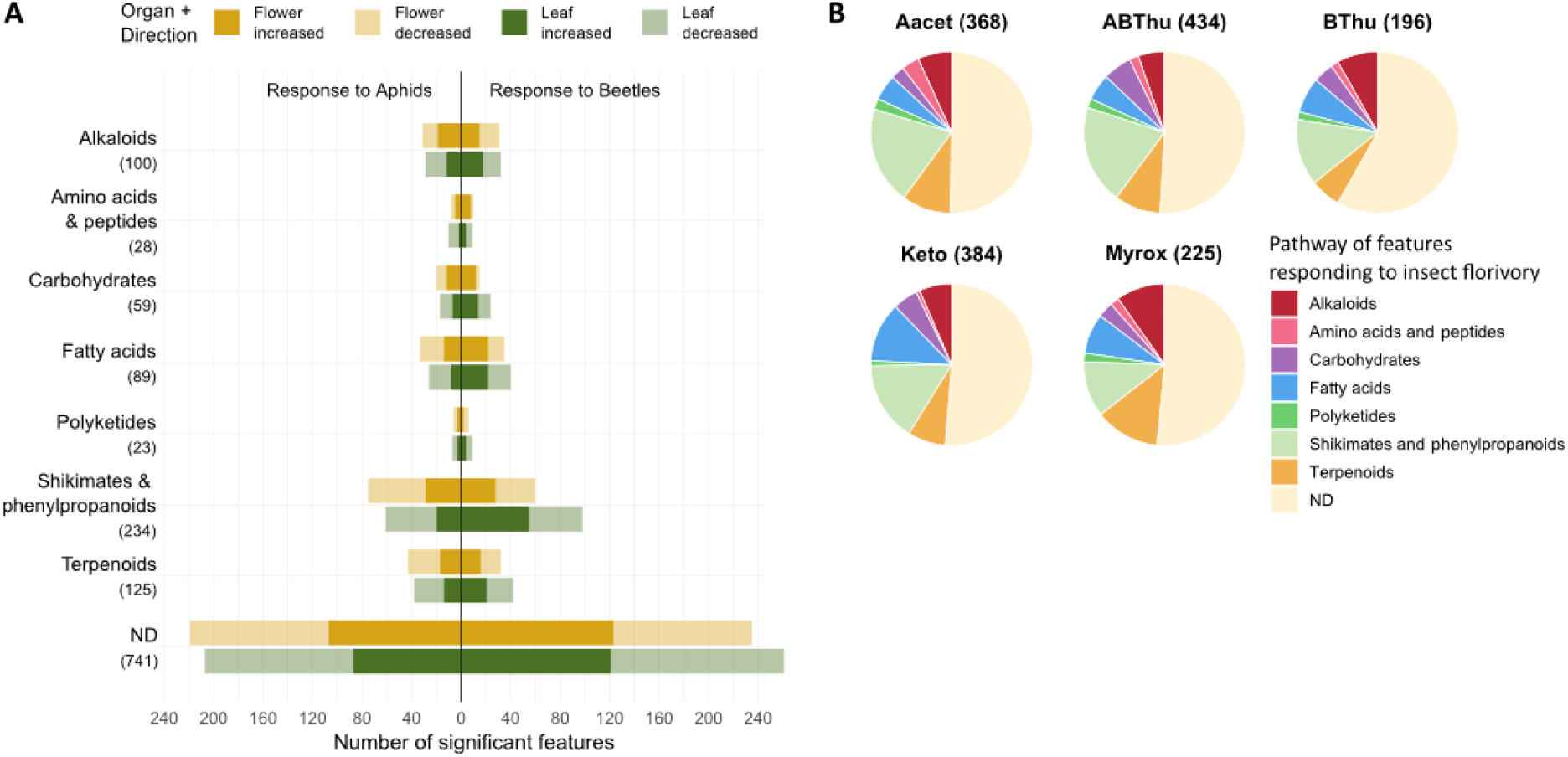
Pathway classification of significant features induced in *Tanacetum vulgare* flower heads and leaves by aphid and beetle florivory. Significant features were identified by moderated t-tests and fold-change analyses conducted using the pooled data set as well as chemotype-specifically (aphid or beetle *versus* control, *p* < 0.05 and fold change < 0.5 for decreases or > 2 for increases). (A) Bar plots showing the number of significant florivory-responsive features, grouped by chemical pathway. The numbers below the pathway labels on the left indicate the total number of unique significant features assigned to each pathway (1399 features in total). For each pathway, the sum of features across the four groups may exceed the total number of significant features assigned to that pathway, because a single feature can be significant in more than one group. ND = not determined. (B) Pie charts showing the distribution of florivory-responsive features across chemical pathways for each chemotype, regardless of organ and insect species. Numbers in parentheses indicate the total number of responding features within each chemotype. Chemotypes are abbreviated as follows: Aacet, artemisyl acetate/artemisia ketone/artemisia alcohol mix chemotype; ABThu, α-/β-thujone mix chemotype; BThu, β-thujone chemotype; Keto, artemisia ketone chemotype; and Myrox, (*Z*)-myroxide/santolina-triene/artemisyl acetate mix chemotype.

### 3.4 Overall Comparison of Florivore-Induced Responses across Analytical Platforms

Of the 159 stored terpenes detected in the flower heads, 96 were also emitted (VOCs). Of these 96 compounds, 21 responded to florivory in the VOCs and 10 in the stored fraction, but only two unknown terpenes (RI1080 and RI1296) responded in both fractions. Across all three analytical platforms, the overlap of metabolites responding to the two insect species was low (Figure 6), meaning that most responding features were specific to one insect species. Moreover, only few features responded in flower heads and leaves simultaneously, in both stored terpenes and metabolic fingerprints, indicating an organ-specific response. Features that responded significantly in only one chemotype clearly outnumbered those that were significant in two or more chemotypes, highlighting chemotype-specific responsiveness.

**FIGURE 6.**
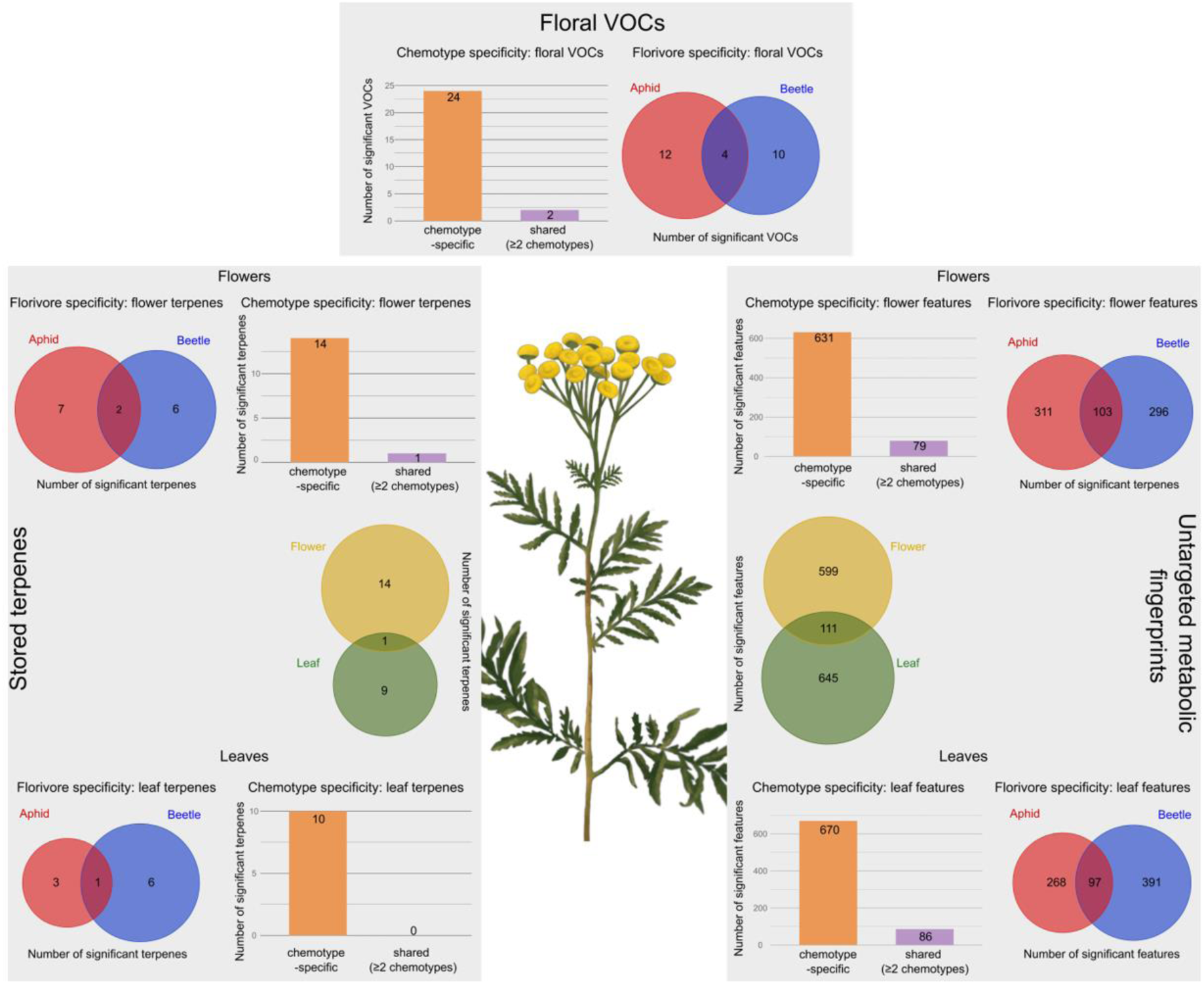
Overall comparison of florivore-induced metabolic responses across three analytical platforms: floral VOCs (TD-GC-MS), stored terpenes (GC-MS) and untargeted metabolomic fingerprints (UHPLC-QTOF-MS/MS). The figure summarises numbers of significant metabolites in *Tanacetum vulgare* floral volatiles, flower heads and leaves following aphid or beetle florivory. Significant metabolites were identified separately for each chemotype using moderated t-tests and fold-change analyses (aphid or beetle *versus* control, *p* < 0.05 and fold change < 0.5 for decreases or > 2 for increases). Venn diagrams show metabolites responding to aphid or beetle florivory or differences among flower heads and leaves, with shared metabolites presented in the intersections.

## 4. Discussion

Our knowledge of induced plant responses to insects with different feeding modes is largely based on leaves. In contrast, floral responses to different types of florivory and their systemic effects on leaf chemistry, have received less attention. Moreover, little is known about how intraspecific diversity in chemotypes affects these responses. Here, we investigated how feeding by florivores differing in feeding mode shapes the metabolic responses of flower heads and leaves in different *T. vulgare* chemotypes. Constitutive defences (total terpene content) were much higher in flower heads than in leaves, while overall terpene profiles of flower heads and leaves remained largely stable after infestation. Nevertheless, both local and systemic responses occurred in individual metabolites in an organ-specific manner. The responding features were mainly related to the shikimate and phenylpropanoid pathways, or were minor terpenes. The number and identity of responding features also differed among chemotypes and depended on the insect species, indicating that the chemotype modulates how strongly plants respond to different flower infestation.

### 4.1 Stable Overall Profiles with Selective, Organ-Specific Responses to Florivory

The overall volatile and stored metabolic profiles of flower heads were not altered by florivory. This is in line with our expectations: flower heads already exhibit high constitutive defences and may therefore require little induction upon attack (Kessler and Halitschke, 2009; McCall and Irwin, 2006). Consequently, the overall profile of leaves, which could respond via systemic signalling, was also largely unaffected. Maintaining a relatively stable floral chemical profile is likely beneficial, as extensive changes in floral chemistry may interfere with pollinator attraction. Recent evidence suggests that floral scent emission can remain remarkably stable following florivory across different plant species: in *Centrosema pubescens* (Fabaceae) and *Tocoyena formosa* (Rubiaceae), florivory did not alter the overall scent profile and only a few volatiles responded (Tunes et al., 2026). In *T. vulgare,* individual metabolites changed in abundance in flower heads, but additionally in leaves. The vascular connection between flowers and leaves may explain such systemic responses. Furthermore, wound-induced signals such as electrical signals and calcium waves, as well as volatiles released from damaged tissues, can coordinate responses in distant tissues and thereby mediate within-plant communication (Heil and Karban, 2010; Lee and Seo, 2022; Orians, 2005). Across all chemotype × insect × organ combinations, more of the responding features decreased rather than increased upon florivory, suggesting a selective increase of a few compounds and an adjusted metabolism. Moreover, these responses were largely organ-specific, while only a comparatively small number responded in both organs. Similar patterns have been observed after leaf herbivory, where some compounds were induced in both flowers and leaves, while others were induced in only one organ in *Solanum peruvianum* (Solanaceae) (Kessler and Halitschke, 2009). This suggests that a common wound signal can trigger organ-specific metabolic responses. It may also reflect the different ecological functions of reproductive and vegetative tissues, as floral chemistry is involved in both pollinator attraction and defence against florivores, whereas leaf chemistry primarily mediates interactions with herbivores and their natural enemies (Kessler and Chautá, 2020; Mithöfer and Boland, 2012).

### 4.2 Stored and Emitted Terpenes are Only Partly Overlapping

Stored and emitted terpenes were only partly matching in *T. vulgare*. Both fractions shared the major terpenes and the leaf and flower chemotypes were reflected in the floral VOCs. Upon infestation, the chemotype-defining terpenes were only weakly altered, while responses were mainly restricted to minor compounds. However, the volatile and stored fractions differed in their responsiveness, with only two terpenes responding in both pools. The different responsiveness may be due to the fact that VOCs were sampled one day earlier than the flower heads and leaves for analysis of stored terpenes due to logistical reasons. A close connection between the volatile and stored fractions has been demonstrated in several *Chrysanthemum* species (Asteraceae), where floral terpene emission rates largely correlated with stored concentrations in undamaged plants (Zhang et al., 2021). However, to our knowledge, stored and emitted terpenes have not been compared in flowers after florivory. In leaves, emissions increased pronouncedly in *Melaleuca alternifolia* (Myrtaceae) after feeding by *Paropsisterna tigrina* beetles and the VOC composition reflected both the constitutive terpene profile and compound volatility, indicating that stored terpenes are the main emission source (Bustos-Segura and Foley, 2018). In contrast, in *T. vulgare* leaves, constitutively stored and emitted terpene profiles only partly overlapped, as some stored terpenes are poorly volatile (Clancy et al., 2016). Responses to herbivory were likewise decoupled between pools in *T. vulgare* shoots, where phloem-feeding aphids increased monoterpenoid emissions without affecting stored fractions (Newrzella et al., 2026).

Similarly, herbivory of *Spodoptera exigua* increased total volatile emissions of *Gossypium hirsutum* (Malvaceae) leaves but hardly changed stored mono- and sesquiterpenes and some compounds, such as α-pinene, were induced in the headspace but not in leaf tissue (Mamin et al., 2025). Chemotype-characteristic terpenes were induced only weakly, as in our data. The terpenes are usually stored in glandular trichomes, while the emission of VOCs highly depends on the morphology of the trichomes and the type and physicochemical properties of the compounds, but the transport processes of VOCs are still not fully understood (Tissier et al., 2017). Thus, VOC and stored terpene profiles may not necessarily overlap, neither under constitutive nor under induced conditions.

### 4.3 Florivore- and Chemotype-Specific Responses to Florivory

Beetles and aphids, applied to separate clones in the laboratory, altered the flower head metabolome of *T. vulgare* to a similar extent in the present study. In contrast, when aphids and beetles were applied to separate stems of the same plant in the field, aphids had a weaker effect than the beetles (Gaar et al., 2026). Systemic signalling within the plant and variable environmental conditions may have led to these different outcomes. Here, we found that *T. vulgare* leaf response was insect species-specific, with beetles inducing slightly more systemic changes than aphids. Similarly, chewing caterpillars of *Spodoptera littoralis* systemically induced terpenes in undamaged leaves of *G. hirsutum*, whereas phloem-feeding aphids *Aphis gossypii* did not systemically affect specialised metabolites (Eisenring et al., 2018). Our results show that the nature of the responsive metabolites was also insect species-specific, which corroborates previous observations (Gaar et al., 2026). In *T. vulgare*, aphid infestation predominantly decreased metabolite levels, while beetles tended to cause an increase (Gaar et al., 2026). This contrasting pattern was only observed in the Aacet chemotype in this study, showing the need for further investigation on floral responses to florivores with different feeding modes. Chewing and phloem-feeding insects can elicit distinct chemical responses. In leaves and other vegetative organs, aphids and chewing herbivores induced different volatile blends in *Solanum tuberosum* (Gosset et al., 2009) and differentially regulated glucosinolates in young *Brassica rapa* (Sotelo et al., 2014). Feeding-type-specific effects also extend to flowers when herbivores attack leaves, as shown in two Brassicaceae species. In *Sinapis alba*, damage by the aphid *Lipaphis erysimi* on vegetative plant parts reduced floral VOC emission, while the chewing caterpillar *Plutella xylostella* had no effect (Pareja et al., 2012). Such insect-specific metabolic responses may reflect the insects’ contrasting feeding modes and the signalling pathways they trigger (Koornneef and Pieterse, 2008; Pieterse et al., 2012), as in *G. hirsutum*, where caterpillars increased jasmonoyl-L-isoleucine and abscisic acid levels, whereas aphids decreased salicylic acid levels (Eisenring et al., 2018).

Plant responses to florivory were not only florivore-specific, but also chemotype-specific, as indicated above for the distinct responses in the Aacet chemotype. Moreover, only a few responding terpenes and untargeted features were shared between chemotypes and the number of responding features varied among chemotypes depending on the florivore. These results are also congruent with previous observations (Gaar et al., 2026). Chemotype-dependent induction has also been observed in other organs of *T. vulgare*. Root-chewing wireworms (*Agriotes* spp.) induced variable responses of stored root sesquiterpenoids between *T. vulgare* chemotypes, while aphids (*M. tanacetaria*) increased shoot monoterpenoid emissions in only one chemotype (Newrzella et al., 2026). In *G. hirsutum*, leaf emissions of aldoximes and nitriles, as well as stored concentrations of (*E*)-β-ocimene, varied between chemotypes after caterpillar feeding (Mamin et al., 2025) and terpene emissions following artificial damage differed among chemotypes of *M. alternifolia* (Bustos-Segura and Foley, 2018). Chemotype-specific responses of terpenes to florivory may be linked to differences in constitutive terpene profiles and the underlying biosynthetic pathways, possibly through chemotype-specific regulation of terpene synthase genes, as proposed for *T. vulgare* roots and shoots (Newrzella et al., 2026). However, chemotype-specific responses were also observed for other, non-terpenoid features, indicating that these responses extend beyond terpene metabolism. Indeed, terpene chemotypes of *T. vulgare* also differ in their broader constitutive metabolome (Dussarrat et al., 2023), while alkaloids and fatty acids allowed chemotype prediction with over 70% accuracy (Müller et al., 2026). Chemotype-specific volatile responses may also act as within-plant signals and thereby modulate induced responses in other metabolic pathways, including non-volatile features (Frost et al., 2007; Heil and Karban, 2010).

### 4.4 Shared Metabolic Pathways behind Specific Responses to Florivory

Although the identity of the responding metabolic features was insect species- and chemotype-specific, most annotated features were associated with the shikimate and phenylpropanoid pathways, as also found in *T. vulgare* flower heads in an earlier study (Gaar et al., 2026). These pathways produce numerous defence-related metabolites, including caffeic acid derivatives, flavonoids and other phenolic compounds, which can act as feeding deterrents, antioxidants or toxins and are frequently induced upon herbivory (Mostafa et al., 2022; War et al., 2012). Many responding features were also annotated as terpenoids and alkaloids in the present study. The responsiveness of terpenoids is consistent with their central role in plant-insect interactions in *T. vulgare*. Alkaloids are also well-known defence compounds (Mostafa et al., 2022; War et al., 2012), that can accumulate following herbivore attack sometimes together with phenolic defences (Fernandes et al., 2011). As alkaloids also differ between *T. vulgare* chemotypes (Müller et al., 2026), they may contribute to the chemotype-specific responses observed here. A few amino acids and peptides responded to florivory in contrasting patterns, with mostly increased in flower heads and decreases in leaves. Although based on few compounds, this tendency may hint at a shift in nitrogen towards the flower heads, which are important nitrogen sinks that can receive nitrogen remobilised from vegetative tissues (Tegeder and Masclaux-Daubresse, 2018).

## 5. Conclusion

In conclusion, the overall metabolic profiles of flower heads and leaves remained largely stable following florivory. Nevertheless, feeding by both florivore species triggered local responses in flower heads and systemic responses in leaves. The magnitude and composition of these responses depended on the florivore’s feeding mode, plant chemotype and organ. This pattern was consistently observed across floral VOC emissions and stored terpenes and untargeted metabolic fingerprints of flower heads and leaves. Despite the differences in individual metabolic responses, the same major compound classes were affected by both florivores, particularly features associated with the shikimate and phenylpropanoid pathways, as well as terpenes, across both organs and all chemotypes. Taken together, these findings show that plant chemistry is shaped by the interplay of florivore identity and intraspecific chemodiversity. Future work could build on this by testing whether such metabolic shifts have ecological consequences, for example for pollinator attraction, under natural conditions, where plants face diverse insect communities and variable environmental conditions.

## Acknowledgements

We acknowledge support for the publication costs by the Open Access Publication Fund of Bielefeld University. We thank Annika Schindéle for her help with plant maintenance.

## Conflicts of Interest

The authors declare no conflicts of interest.

## Data Availability Statement

Chemical data, metadata and code will be made available upon acceptance of this manuscript.

## Funding

This research was funded by the German Research Foundation (Deutsche Forschungsgemeinschaft, DFG), project MU1829/28-2 and DU 2440/1-2, as part of the research unit FOR 3000 (project number 415496540).

## Supporting Information

Additional supporting information can be found online in the Supporting Information section

## Notes

### Competing Interest Statement

The authors have declared no competing interest.

