## Supplementary Figures for "Effects of Florivore Type and Plant Chemotype on Metabolic Responses in Flowers and Leaves of an Aromatic Plant Species"

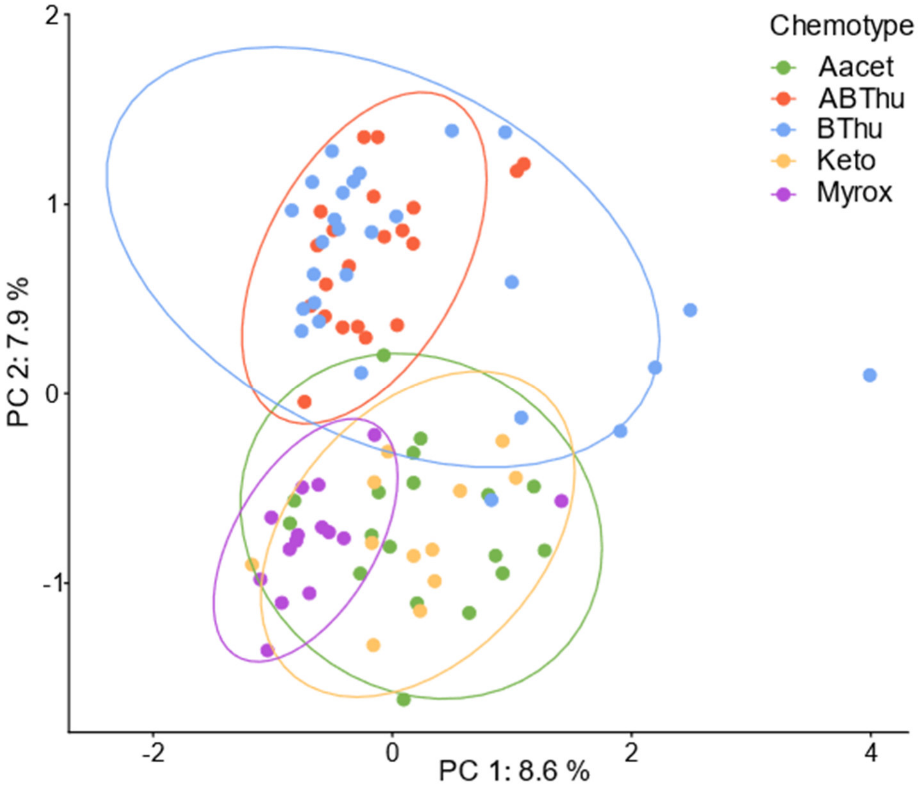


**Fig. S1. PCA of volatile profiles of flower heads of all *Tanacetum vulgare* samples belonging to different chemotypes.** Chemotypes are dominated by artemisyl acetate, artemisia ketone and artemisia alcohol (Aacet); α-thujone and β-thujone (ABThu); β-thujone (BThu); artemisia ketone (Keto); and (*Z*)-myroxide, santolina triene and artemisyl acetate (Myrox). PCA score plot, based on normalised TD-GC-MS data of 136 volatile organic compounds (VOCs), shows a clear separation between chemotypes.

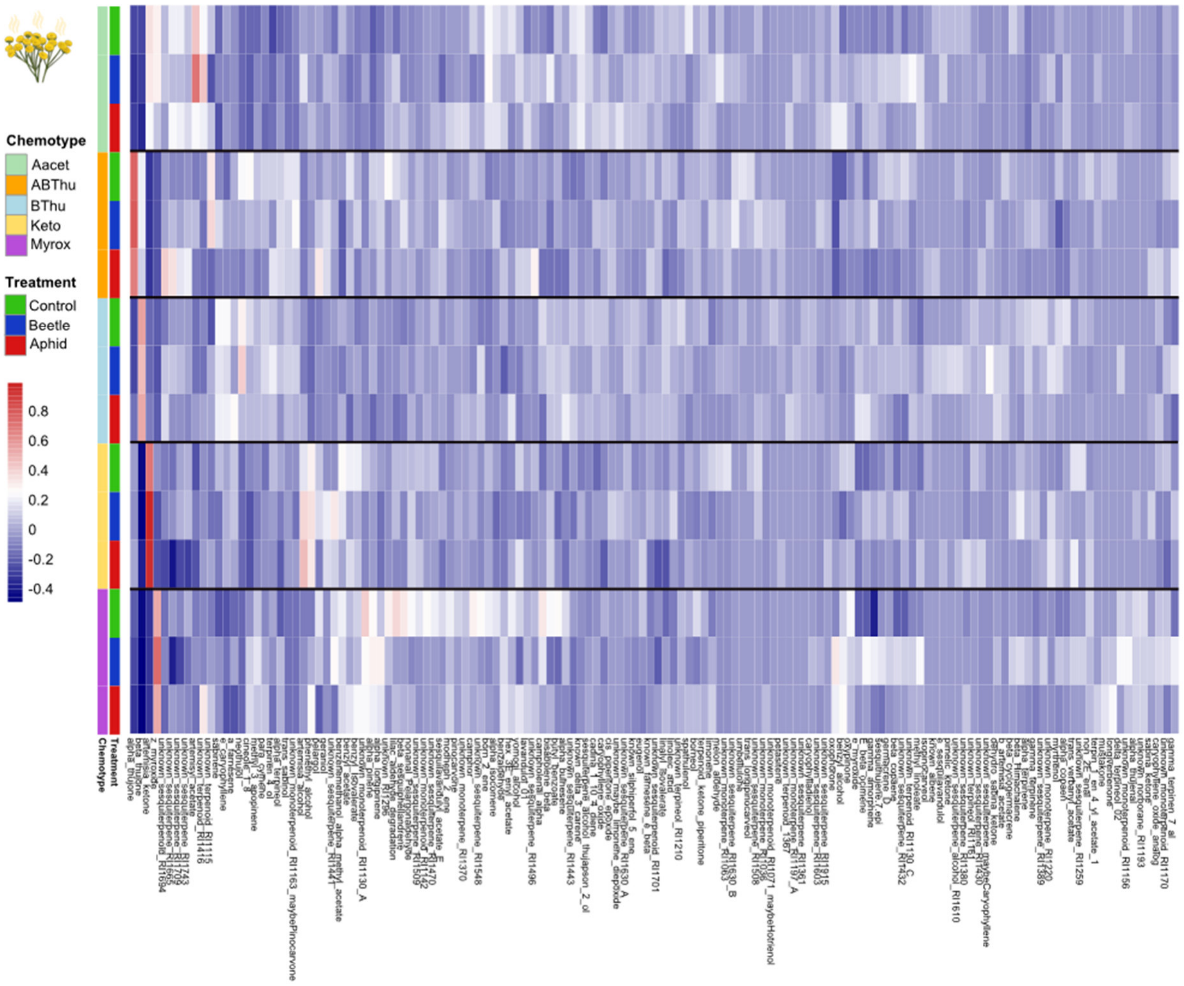


**Fig. S2. Heatmap of the group mean values of normalised abundances of all floral volatile organic compounds (VOCs) detected in *Tanacetum vulgare*.** Rows correspond to treatment groups (control: uninfested; beetle: infested by *Olibrus* spp.; aphid: infested by *Macrosiphoniella tanacetaria*) within each chemotype. Columns correspond to the VOCs, clustered based on Pearson correlation and colours of different intensities represent normalised abundances. Chemotypes are abbreviated as follows: Aacet, artemisyl acetate/artemisia ketone/artemisia alcohol mix chemotype; ABThu, α-/β-thujone mix chemotype; BThu, β-thujone chemotype; Keto, artemisia ketone chemotype; and Myrox, (*Z*)-myroxide/santolina-triene/artemisyl acetate mix chemotype.

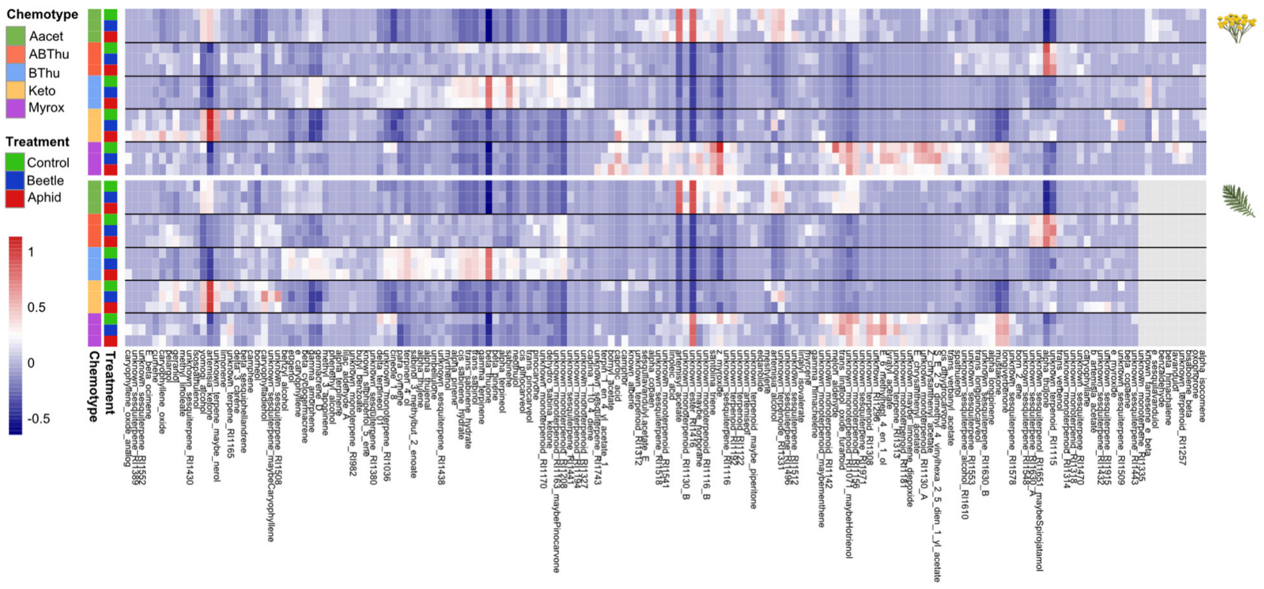


**Fig. S3. Heatmaps of the group mean values of normalised abundances of all stored terpenes in flower heads (top) and leaves (bottom) detected in *Tanacetum vulgare*.** Rows correspond to treatment groups (control: uninfested; beetle: infested by *Olibrus* spp.; aphid: infested by *Macrosiphoniella tanacetaria*) within each chemotype. Columns correspond to the individual terpenes, clustered based on Pearson correlation and colours of different intensities within the heatmaps represent normalised abundances. Grey cells indicate that the respective terpene was not detected in that sample. Chemotypes are abbreviated as follows: Aacet, artemisyl acetate/artemisia ketone/artemisia alcohol mix chemotype; ABThu, α-/β-thujone mix chemotype; BThu, β-thujone chemotype; Keto, artemisia ketone chemotype; and Myrox, (*Z*)-myroxide/santolina-triene/artemisyl acetate mix chemotype.

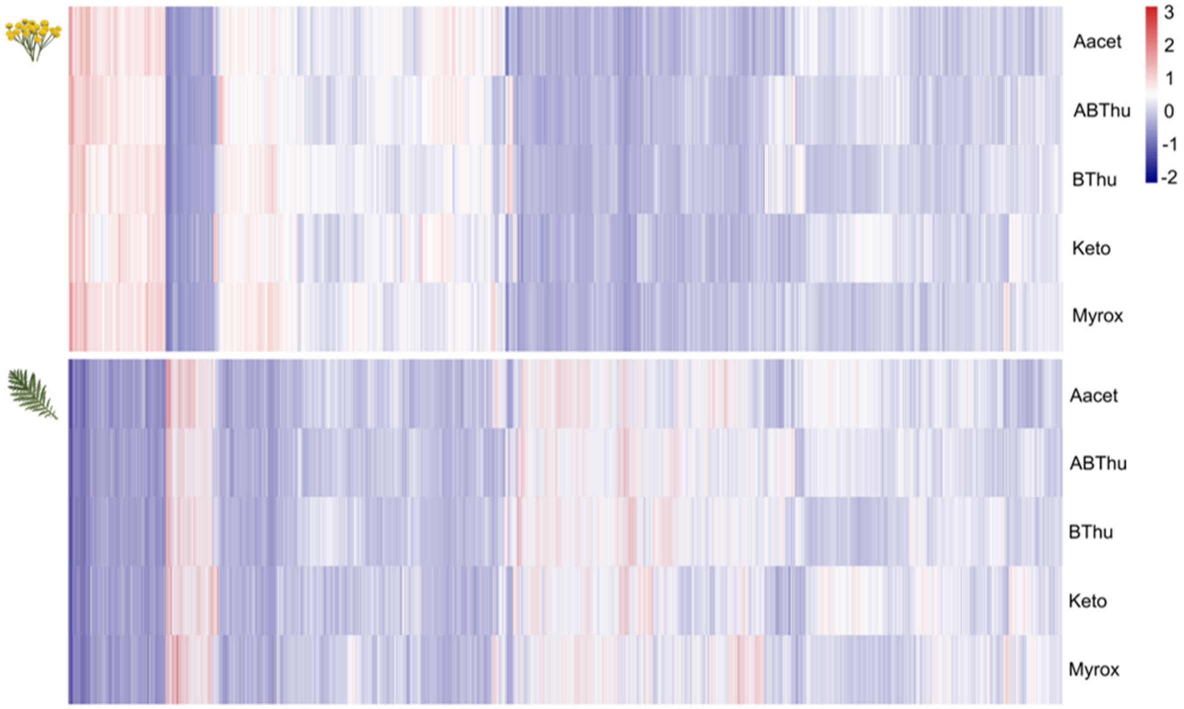


**Fig. S4. Heatmaps of the group mean values of normalised intensities of LC-MS features shared between flower heads (top) and leaves (bottom) in *Tanacetum vulgare* (7122 features across all treatments).** Rows correspond to the chemotype, columns correspond to the individual features, clustered based on Pearson correlation and colours of different intensities within the heatmaps represent normalised abundances. Chemotypes are abbreviated as follows: Aacet, artemisyl acetate/artemisia ketone/artemisia alcohol mix chemotype; ABThu, α-/β-thujone mix chemotype; BThu, β-thujone chemotype; Keto, artemisia ketone chemotype; and Myrox, (*Z*)-myroxide/santolina-triene/artemisyl acetate mix chemotype.

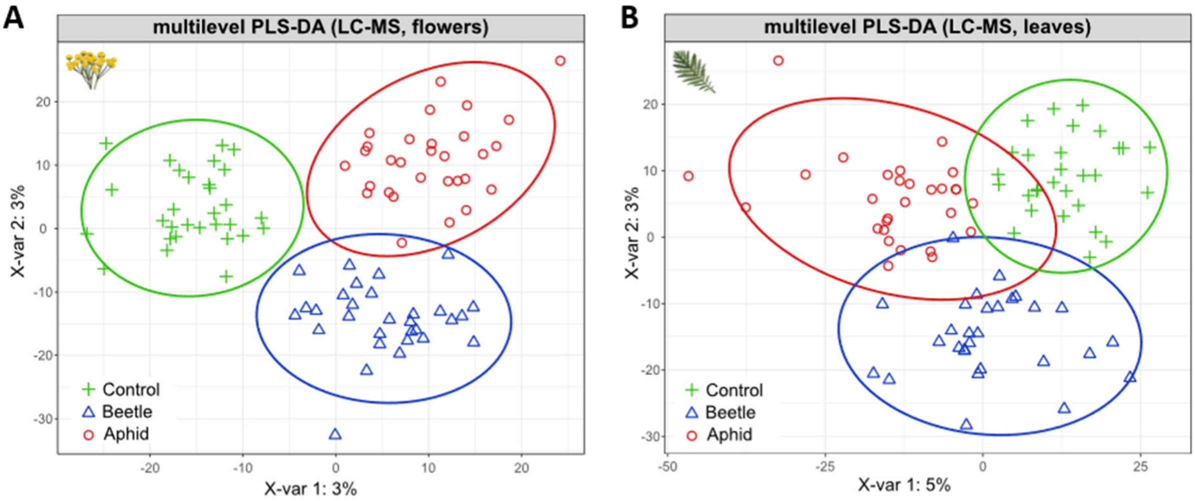


Error rate per class:

| Treatment | Comp 1 | Comp 2 |
|---|---|---|
| Control | 0.237 | 0.305 |
| Beetle | 0.783 | 0.362 |
| Aphid | 0.541 | 0.393 |

Error rate per class:

| Treatment | Comp 1 | Comp 2 |
|---|---|---|
| Control | 0.422 | 0.415 |
| Beetle | 0.835 | 0.427 |
| Aphid | 0.456 | 0.456 |

**Fig. S5. Metabolic fingerprints of (A) flower heads and (B) leaves of cloned *Tanacetum vulgare* plants belonging to five chemotypes in response to insect infestation.** Multilevel PLS-DA score plots of LC-MS metabolic fingerprints illustrating treatment-related variation in the paired and normalised data, comprising 7163 features in flower heads and 7134 features in leaves. Symbols represent treatment groups (control: uninfested; beetle: infested by *Olibrus* spp.; aphid: infested by *Macrosiphoniella tanacetaria*). Ellipses denote 95% confidence intervals for each treatment group. Bottom tables summarise classification error rates for components 1 and 2, reflecting model performance per treatment class.

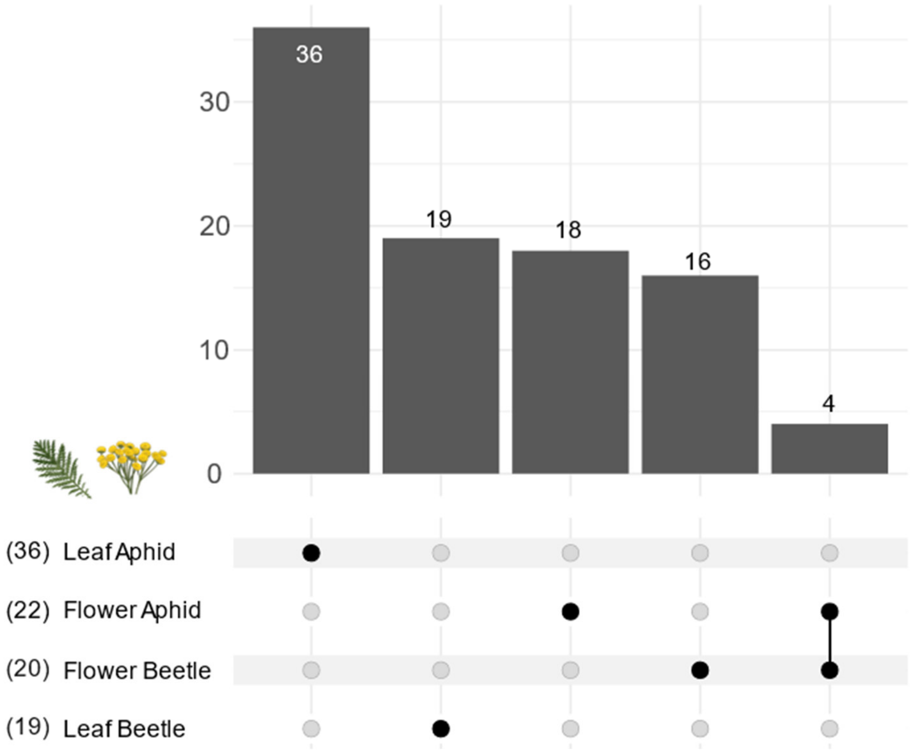


**Fig. S6. UpSet plot of LC-MS features significantly responsive to florivory across flower heads and leaves of *Tanacetum vulgare*, irrespective of chemotype**. Features were identified as significantly altered by moderated t-tests and fold-change analyses for the aphid or beetle treatment *versus* the unifested control ($P < 0.05$; fold change < 0.5 for decreases or > 2 for increases). Numbers in parentheses on the left indicate the total number of significant features within each group, while bars on the top represent intersections between groups, illustrating shared and unique feature responses to insect infestation.
